# Singe cell oil (SCO) based bioactive compounds: II. Enzymatic synthesis of glucose fatty acid esters using SCOs as acyl group-donors and their biological activities

**DOI:** 10.1101/2020.10.30.362673

**Authors:** Hatim A. El-Baz, Ahmed M. Elazzazy, Tamer S. Saleh, Marianna Dourou, Jazem A. Mahyoub, Mohammed N. Baeshen, Hekmat R. Madian, George Aggelis

## Abstract

Sugar fatty acid esters, especially glucose fatty acid esters (GEs), have broad applications in food, cosmetic and pharmaceutical industries. In this research the fatty acid moieties derived from polyunsaturated fatty acid containing single cell oils (SCOs), i.e. those produced from *Cunninghamella echinulata*, *Umbelopsis isabellina* and *Nannochloropsis gaditana* as well as from olive oil and an eicosapentaenoic acid (EPA) concentrate were converted into GEs by enzymatic synthesis, using lipases as biocatalysts. The GE synthesis was monitored using thin-layer chromatography, FT-IR and in situ NMR. It was found that GE synthesis carried out using immobilized *Candida antarctica* B lipase was very effective reaching high yields, near to 100%. It was shown that EPA-GEs were very effective against several pathogenic bacteria and their activity can be attributed to their high EPA content. Furthermore, *C. echinulata-GEs* were more effective against pathogens comparing to *U. isabellina-GEs*, probably due to the presence of gamma linolenic acid (GLA) in the lipids of *C. echinulate*, which is known for its antimicrobial activity, in higher concentrations. *C. echinulata-GEs* also showed a strong insecticidal activity against *Aedes aegypti* larvae, followed by EPA-GEs, olive oil-GEs, and *N. gaditana-GEs*. All synthesized GEs induced apoptosis of the SKOV-3 ovarian cancer cell line, with the apoptotic rate increasing significantly after 48 h. A higher percentage of apoptosis was observed in the cells treated with EPA-GEs, followed by *C. echinulata-GEs, U. isabellina-GEs* and olive oil-GEs. We conclude that SCOs can be used in the synthesis of GEs with interesting biological properties.

## 1. Introduction

Sugar fatty acid esters, so-called sugar esters (SEs), are biodegradable, odorless, non-irritating and non-toxic surfactants with broad applications in food, cosmetic and pharmaceutical industries [1–4]. Moreover, SEs have gained attention thanks to their anti-bacterial (e.g. against numerous pathogenic species of gram-positive and gram-negative bacteria) and anti-fungal activity [5], while they are also reported as insecticide and miticide [6].

SEs are synthesized from renewable resources, such as sugars and fatty acids (FAs). Different types of sugars (e.g. sucrose, fructose, glucose and lactose) can be used as acyl-acceptors to produce SEs by esterification with FAs or transesterification with FA esters as acyl-donors [7–9]. Glucose, a cheap and broadly available carbohydrate, having only one primary hydroxyl group, which predetermines a highly regioselective synthesis of SEs called glucose fatty acid esters (GEs) [10, 11]. Concerning acyl-donors, polyunsaturated fatty acids (PUFAs) can be used for this purpose, as it is known that PUFAs or PUFA derivatives possess interesting biological activities [12–14]. GEs can be synthesized both chemically and enzymatically [15, 16]. Lately, the main efforts in the processing of GEs have been focused on enzymatic synthesis because this is an advantageous method, requiring less energy and reducing solvent toxicity [11, 17], with lipases being the most important enzymes used for this synthesis [18, 19].

Commercially available immobilized lipases, such as *C. antarctica* B lipase, are used to catalyze the acylation of glucose with various FAs [20, 21]. Different sources of PUFAs can be used as sources of acyl groups, in the form of either FAs or FA esters, for GEs production. Whilst the terrestrial plant oils have been used as a source of PUFAs [22, 23], alternative and non-food supply sources, such as microalgae and fungi could be used thanks to their ability to produce microbial lipids rich in PUFAs of medicinal and nutritional interest [24–26]. The microbial lipids, so-called single cell oils (SCOs), are synthesized by oleaginous microorganisms that are capable of producing substantial amounts of lipid stored within their cells [27, 28]. The genus of *Nannochloropsis* includes various marine microalgae species able to efficiently grow under a variety of culture conditions on low quality waters, even on wastewaters, and accumulate lipids rich in PUFAs, such as eicosapentaenoic acid (EPA) [29–32]. Furthermore, oleaginous fungal species, such as *Cunninghamella echinulata* and *Umbelopsis isabellina* are capable to synthesize PUFAs, especially γ-linolenic acid (GLA) [13, 25], thus regarded as promising candidates for SCOs production [33–35].

In a previous paper [36] we produced FA amides (FAAs) using as acyl group-donors lipids containing PUFAs in different percentages and we concluded that FAAs can be used as bioactive compounds in various biological applications depending on their FA composition. The aim of the current paper is to enzymatically synthesize GEs using similar to those used in FAAs SCOs as acyl group-donors, either in the form of free fatty acids (FFAs) or in the form of fatty acid methyl esters (FAMEs). The reaction was carried out under various conditions, using two immobilized lipases as catalysts. SCOs, produced from various sources as describe in El-Baz et al. [36], contained EPA, GLA or oleic acid in high percentages. The biological activity of the aforementioned GEs against important human pathogens, the larvae of *Aedes aegypti* and the SKOV-3 cancer cell line, was studied and compared with GEs synthesized using olive oil and an EPA concentrate (i.e. a fish oil derivative containing EPA in very high percentages) as acyl group-donors. We concluded that GEs derived from SCOs possess interesting biological activities and can therefore be used in the production of pharmaceuticals in the future.

## 2. Materials and methods

### 2.1. Biological material and SCO production

The fungal strains *Cunninghamella echinulata* ATHUM 4411 and *Umbelopsis isabellina* ATHUM 2935 (culture collection of National and Kapodistrian University of Athens, Greece) and the microalga strain *Nannochloropsis gaditana* (culture collection CCAP 849/5) were used as sources of SCOs. Culture conditions, cell mass harvesting, lipid extraction and purification, FAMEs and FFAs preparation and gas chromatography analysis were described in El-Baz et al. [36].

### 2.2. Enzymatic synthesis of GEs

GEs were synthesized by esterification (or transesterification) of glucose (Acros Organics, Thermo Fisher Scientific, Waltham, MA) served as an acyl acceptor, with FFAs (or FAMEs) served as acyl-group donors. The reaction was performed in 50-mL Erlenmeyer flasks using different FA (or FAME):glucose molar ratios (i.e. 1:1, 1:2 or 1:3). The reactants were dissolved in 25 mL of a solvent mixture consisted of 80% DMSO and 20% tert-amyl alcohol (Sigma Aldrich Co.) in which 1 g of 3 Å molecular sieves was added and the mixture was sonicated for 20 min. The reaction was catalyzed by 0.25 g of immobilized *C. antarctica* B lipase (enzymatic activity □5 Units/mg) or 0.25 g of immobilized *C. rugosa* lipase (enzymatic activity □0.1 Units/mg), both purchased from Sigma Aldrich Co., St. Louis, MO. The flasks were incubated at 50 ± 1 C in a shaking incubator at 100 rpm for 50 h.

After the incubation period the reaction mixture was filtered to remove the molecular sieves and the immobilized lipase and the filtrates were evaporated under reduced pressure to remove the solvent. The reaction residue was separated in ethyl acetate and distilled water (25 mL each). The organic layer, containing the synthesized GEs, was washed with 10 mL saturated aqueous NaCl (Sigma Aldrich Co.), dried over MgSO_4_ (Sigma Aldrich Co.), gravity-filtered and the solvent removed under reduced pressure to get the crude product.

### 2.3. GE analysis

#### 2.3.1. Thin-layer chromatography and FT-IR

Qualitative synthesis of GEs was monitored by thin-layer chromatography (TLC) as described in El-Baz et al. [36] for FAAs. Equally, FT-IR spectra for FAMEs and the GE products were recorded as described in the aforementioned paper. FT-IR spectra were used to detect the formation of the ester carbonyl group confirming thus GE synthesis.

#### 2.3.2. Quantitative determination of the reaction yields through in situ NMR monitoring

The % conversion of FAMEs to GEs was determined during the reaction via in situ NMR monitoring. First, the protons NMR of both reactants individually were assigned and the progress of the reaction was monitored by ^1^H NMR at regular intervals of 10 h. The % conversion was calculated according to the formula:

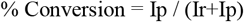

where Ip is the integration of the signal of the product and Ir is the integration of the signal of the reactant.

Heteronuclear multiple bond correlation (HMBC) spectroscopy was used to identify the ester product by determining the chemical shifts of carbon and hydrogen atoms and formation the ester bond. NMR spectra were recorded at 298 K on a Bruker Avance III 400 (9.4 T, 400.13 MHz for 1H, 100.62 MHz for 13C) spectrometer (Bruker, Billerica, MA) with a 5-mm BBFO probe. Chemical shifts (δ in ppm) were relative to internal standard DMSO-d6 (d 2.50) for 1H NMR.

### 2.4. Biological activity of GEs

The antimicrobial, insecticidal and anticancer activities of the synthesized GEs were determined using the protocols described in El-Baz et al. [36]. Briefly, the antimicrobial activity of GEs was evaluated *in vitro* using the agar well diffusion assay [37], the minimum inhibitory concentration (MIC) and minimum bactericidal concentration (MBC) [38] against human pathogens including the Gram-negative *Escherichia coli* ATCC 25922, *Klebsiella pneumoniae* ATCC 700603, *Pseudomonas aeruginosa* ATCC 15442, *Salmonella typhimurium* ATCC 14028 the Grampositive bacteria, *Bacillus subtilis* ATCC 6633, MRSA *Staphylococcus aureus* ATCC 4330, *S. aureus* ATCC 25923 and the unicellular fungus *Candida albicans* ATCC 10221. The insecticidal activity was evaluated by exposing early 4^th^ instar larvae of a field strain of *Aedes aegypti* to different concentrations of the GEs for 48 h, in glass beakers containing 100 mL of tap water. Calculation of statistical parameters was performed using the Finney method [39]. The apoptotic activity of the SKOV-3 ovarian cancer cell line in response to tested compounds (both FAMEs and GEs) was determined by Annexin FITC, as per the manufacturer’s instructions (BD Biosciences, USA).

### 2.5. Statistical analysis

The acquired data were analyzed using SPSS 9.0 and the results were given as mean ± SD of three replicates. The mean comparison between the various assessed groups was performed using one-way analysis of variance (ANOVA). Statistical significance was defined when *p* < 0.05.

## 3. Results and discussion

### 3.1. Biomass and SCO production

The performance of the oleaginous microorganisms cultivated in flasks or bioreactors, as well as the FA composition of the produced lipids, were previously presented [36]. Here, the essential data are provided as supplementary material to facilitate reading (Tables S1, S2).

### 3.2. Optimization of the GE synthesis

For GE synthesis, the olive oil derived FFAs and FAMEs were used as model substrates to optimize the lipase-catalyzed reaction of esterification and transesterification (Fig. 1). The reaction was carried out in a solvent system consisted of tert-amyl alcohol and DMSO mixture as describe elsewhere [11, 40, 41], for 50 h at 55 C with shaking at 100 rpm and the progress of the reaction was monitored by TLC analysis.

**Fig. 1.**
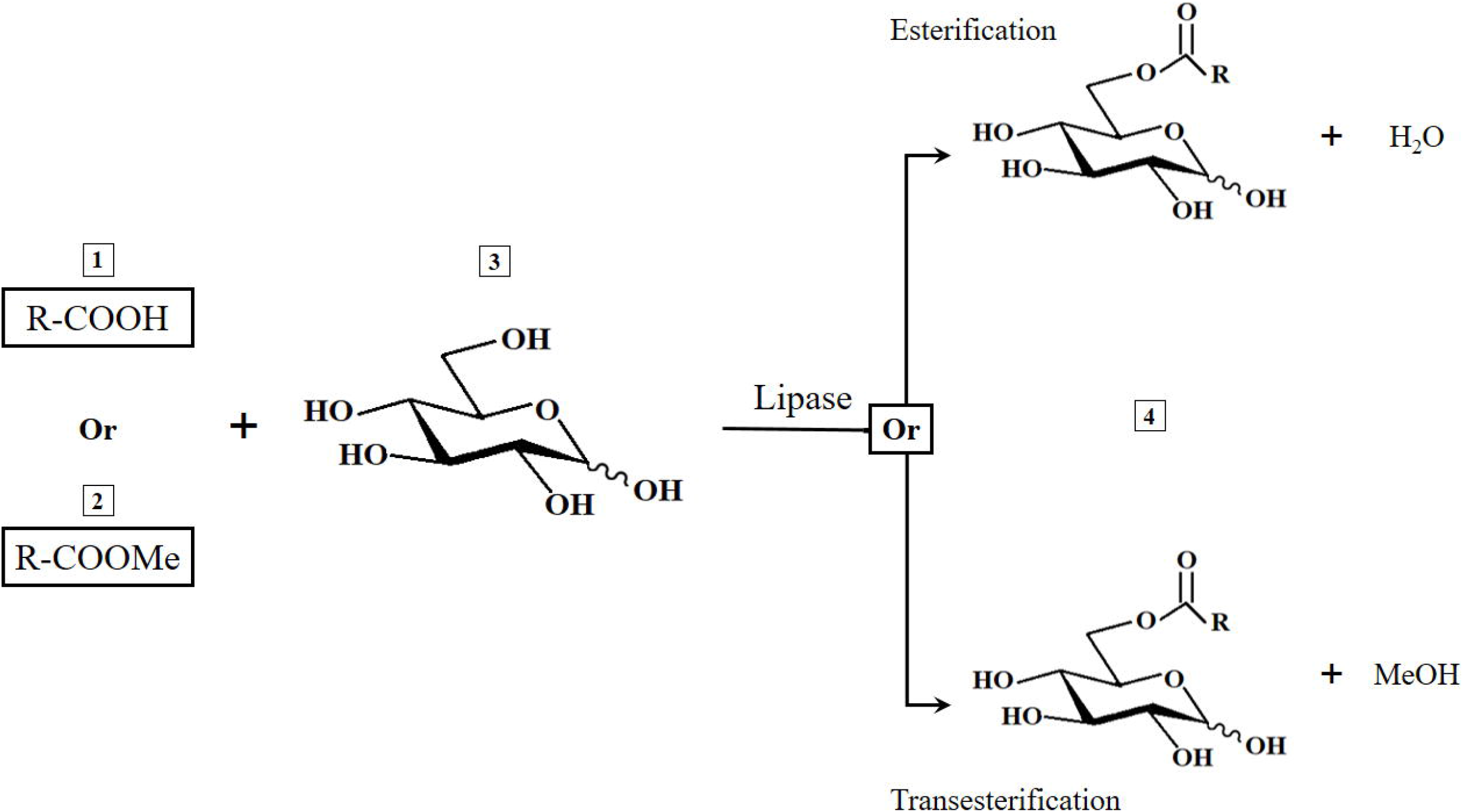
Enzyme-catalyzed synthesis of glucose esters using free fatty acids (FFAs) or fatty acid methyl esters (FAMEs) as substrates.

The effect of various reaction conditions, such as the molar ratio of the reactants, the reaction temperature and the type and the concentration of immobilized enzyme on the conversion rate and yield was studied. Two immobilized lipases, lipase from *C. antarctica* B (Lipase CA) and lipase from *C. rugosa* (Lipase CR) were used as catalysts for both esterification and transesterification reactions for GEs production (Table 1). The employment of immobilized lipases as catalysts for GEs synthesis has been proposed by several researchers [9, 21, 42-44]. Also, different molar ratios of olive oil FFAs (or FAMEs):glucose were tested (Table 1), showing that the conversion yield increased with increasing the concentration of glucose, attaining 100% conversion with lipase CA as catalyst and FAMEs:glucose ratio 1:3 (entry 6, Table 1 and Fig. 2). Glucose can be considered as a good acyl acceptor for SE synthesis in non-conventional media, ensuring high conversion yields, due to its relatively high solubility [45-47]. Contrary, the use of sugars with higher degree of polymerization adversely affects the conversion yield as a result of the low solubility of sugar in organic solvents.

**Fig. 2.**
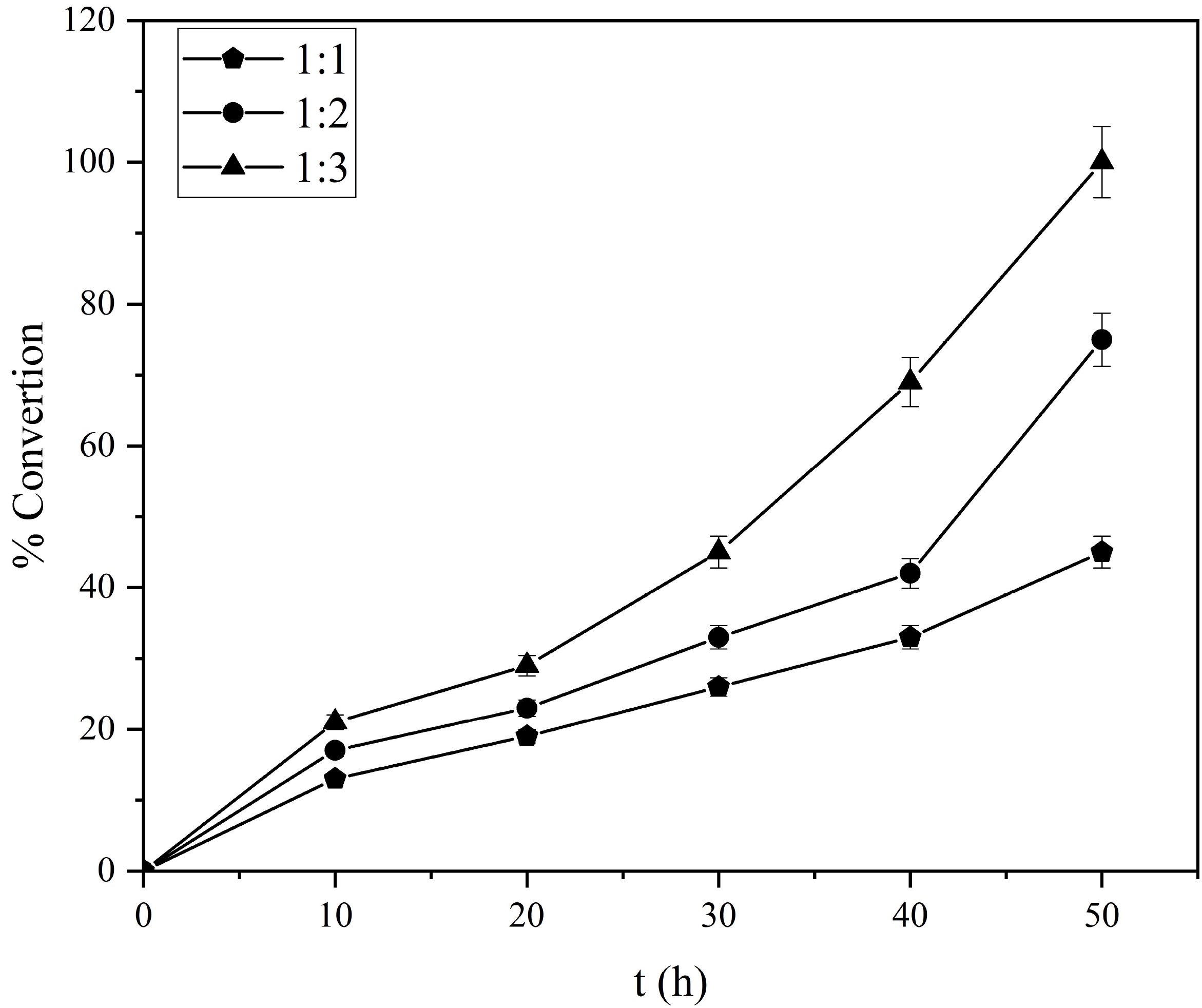
Reaction course over time of olive oil FAMEs conversion to glucose esters in different olive oil FAMEs:glucose ratios.

**Table 1.**
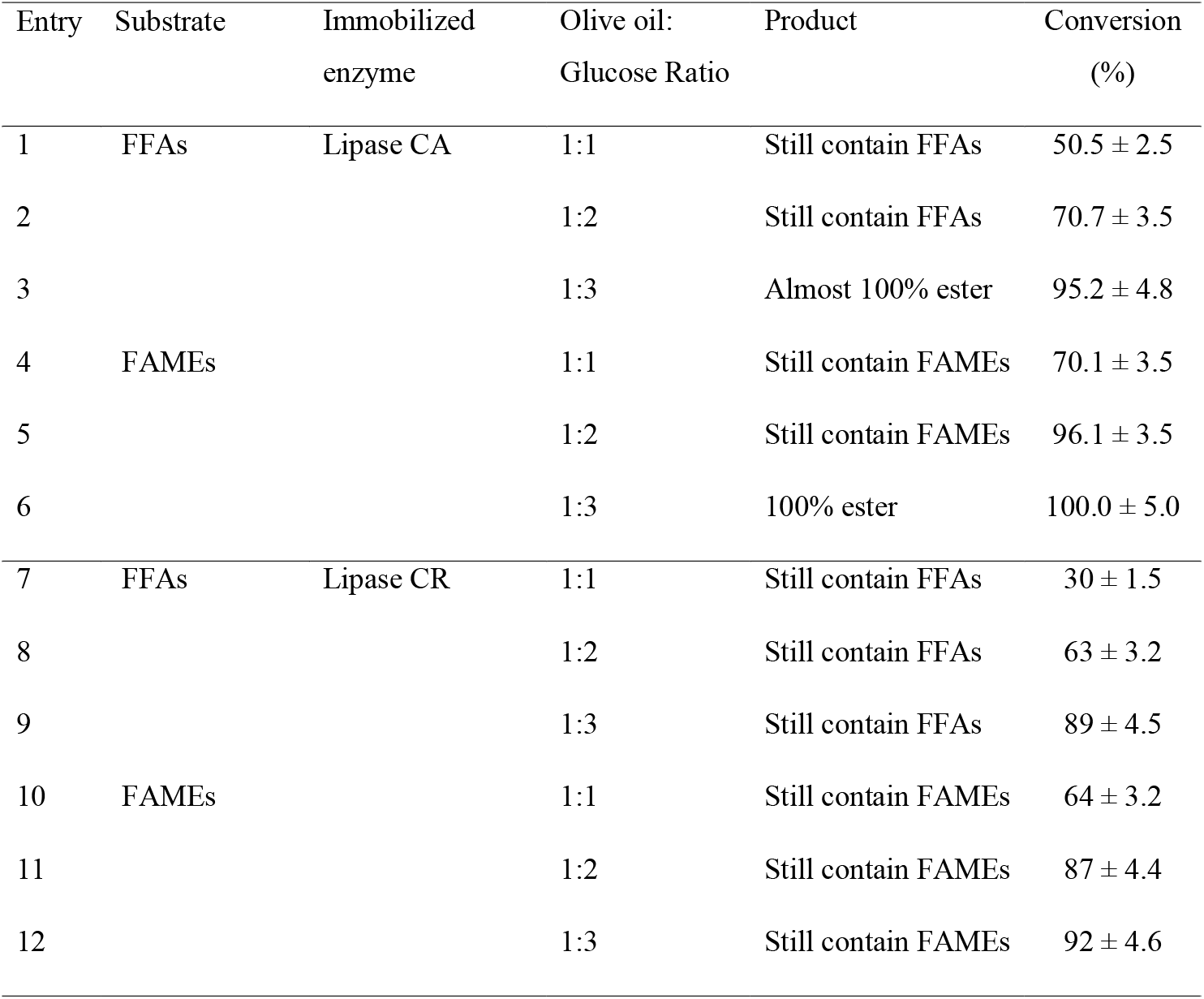
Comparison of GE synthesis using two immobilized lipases and olive oil derived FFAs or FAMEs as substrates at different molar ratio of the reactants.

Although the conversion yield obtained utilizing FFAs as acyl donor was high, this was in all cases below to that obtained with FAMEs as substrate due to the water that was produced, which may react with GE causing hydrolysis (Fig. 1). Contrary, the transesterification reaction carried out using FAMEs is irreversible, with methanol as a byproduct acting as an internal water scavenger, improving thus the conversion yield [48, 49]. Lipase CR showed lower conversion yields (entries 7-12, Table 1) than lipase CA. Furthermore, the reusability of lipase CA was checked for several reaction cycles for the synthesis of GE using olive oil FAMEs as substrate under the optimized reaction conditions. Specifically, the enzyme was removed after the completion of the reaction by filtration, washed with ethanol solvent in a Soxhlet extraction apparatus and reused three times under the same reaction conditions. It was found that the regenerated enzyme performed the reaction efficiently without loss of its catalytic activity, which is important for the sustainability of the process. The reaction conditions were further optimized using the lipase CA in different quantities as a catalyst and it was found that a 100% conversion yield was obtained using 0.25 g of the lipase CA (Table 2, entry 4), while a higher enzyme quantity was not necessary.

**Table 2.**
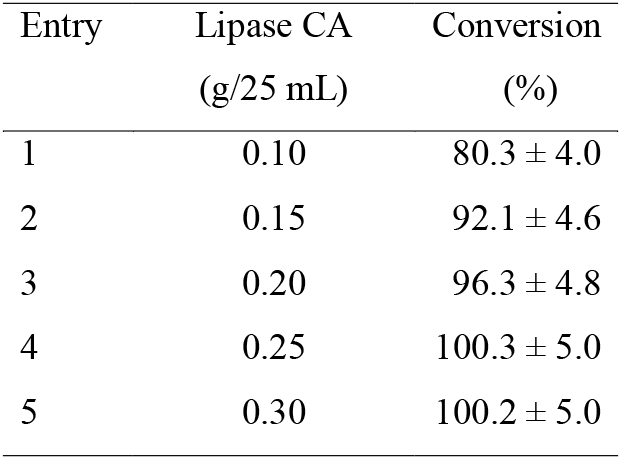
GE synthesis using olive oil derived FAMEs (at a ratio FAMEs:Glc 1:3), utilizing lipase CA as catalyst at different concentrations, in a solvent mixture consisted of 80% DMSO and 20% tert-amyl alcohol and for a duration of incubation 50 h.

The results recorded in this study emphasized the feasibility of enzymatic synthesis of GEs, the conversion yield of which reached 100% using lipase CA as biocatalyst. These results are in agreement with those reported by Yan et al. [50] who demonstrated that glucose FA monoesters were synthesized (with up to 93% yields) using lipase B from *C. antarctica*. High yields for GE enzymatic synthesis were also reported by Sebatini et al. [51] using as catalyst lipase-Fe_3_O_4_ nanoparticles. Furthermore, Findrik et al. [20] reported that the highest SE yield was achieved using lipase CA as catalyst and glucose and palmitic acid as substrates.

### 3.3. Product identification and quantitative analysis

The percent conversion, which was taken as a scale to determine the optimum conditions, was quantified via *in situ* NMR monitoring (Fig. 3). The ^1^H NMR was noted new multiple signals at 4.17, 4.41 ppm, which matched the CH_2_OCO grew concurrently with a decline in the intensity of the glucose protons of C6 signals at 3.39 and 3.52 ppm. The latter signals disappeared after 50 h of reaction when FAMEs:glucose 1:3 ratio was used, which means that 100% conversion was achieved (Fig. 3). The main product obtained was identified as Glc (C-6)-OCOR, as indicated by 2D (^1^H-^13^C HMBC) NMR (Fig. 4a), showing correlation between the peak assigned to proton C-6 glucose ester and the carbonyl function in olive oil FAMEs (Fig. 4b). The structure of the obtained GEs was additionally confirmed by FT-IR analysis showing the appearance of a broadband at 3380 cm^-1^ due to the hydroxyl groups of glucose and the two stretching bands of O-C bond at 1316 and 1015, in parallel with the disappearance of the band at 1743 cm^-1^, due to the consumption of the carbonyl group of FAMEs (Fig. 5).

**Fig. 3.**
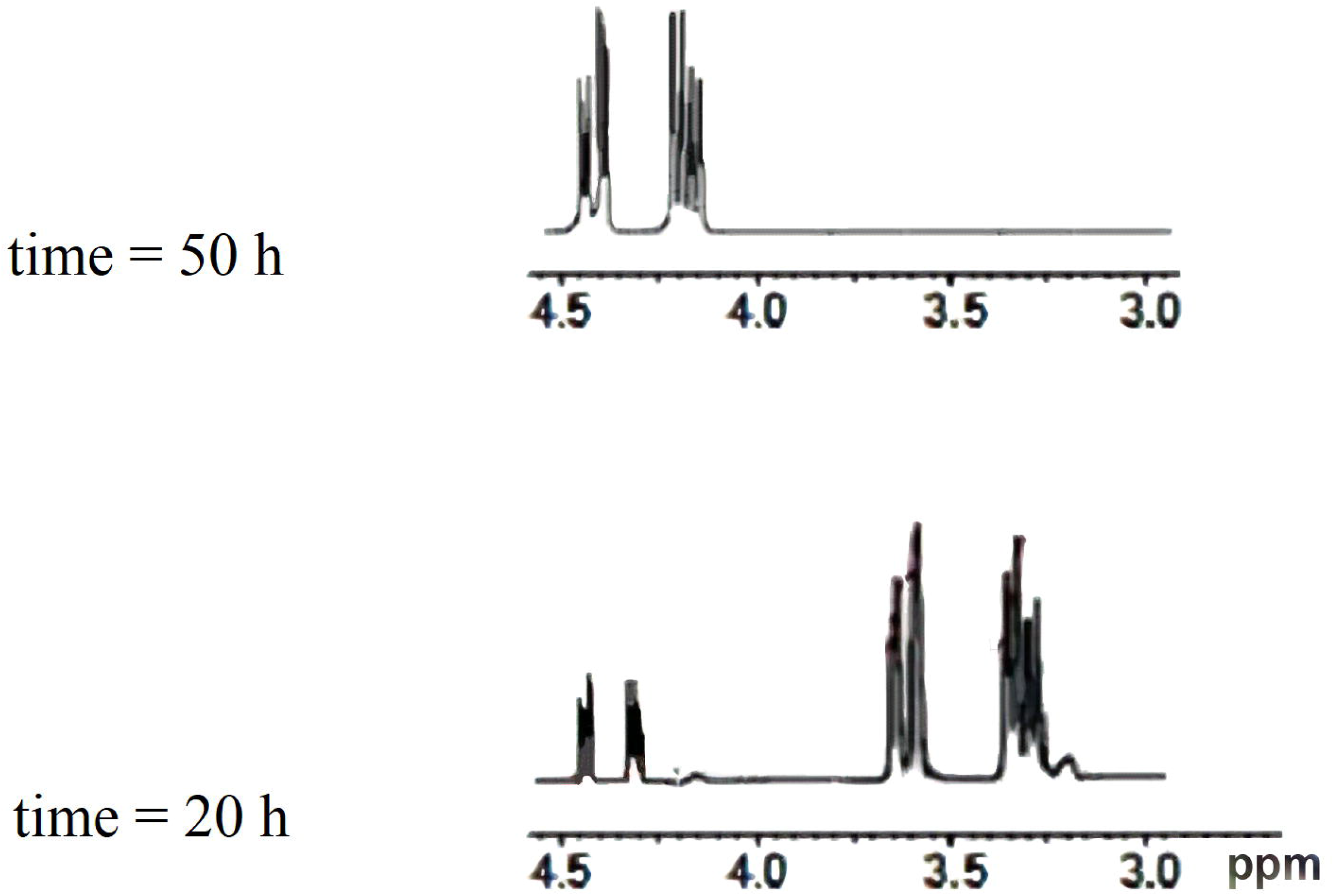
NMR monitoring of glucose ester synthesis. The signal intensity at 4.17 and 4.41 ppm derived from the CH_2_OCO group increased simultaneously with a decrease of the signal intensity derived from the glucose protons of C6 at 3.39 and 3.52 ppm.

**Fig. 4.**
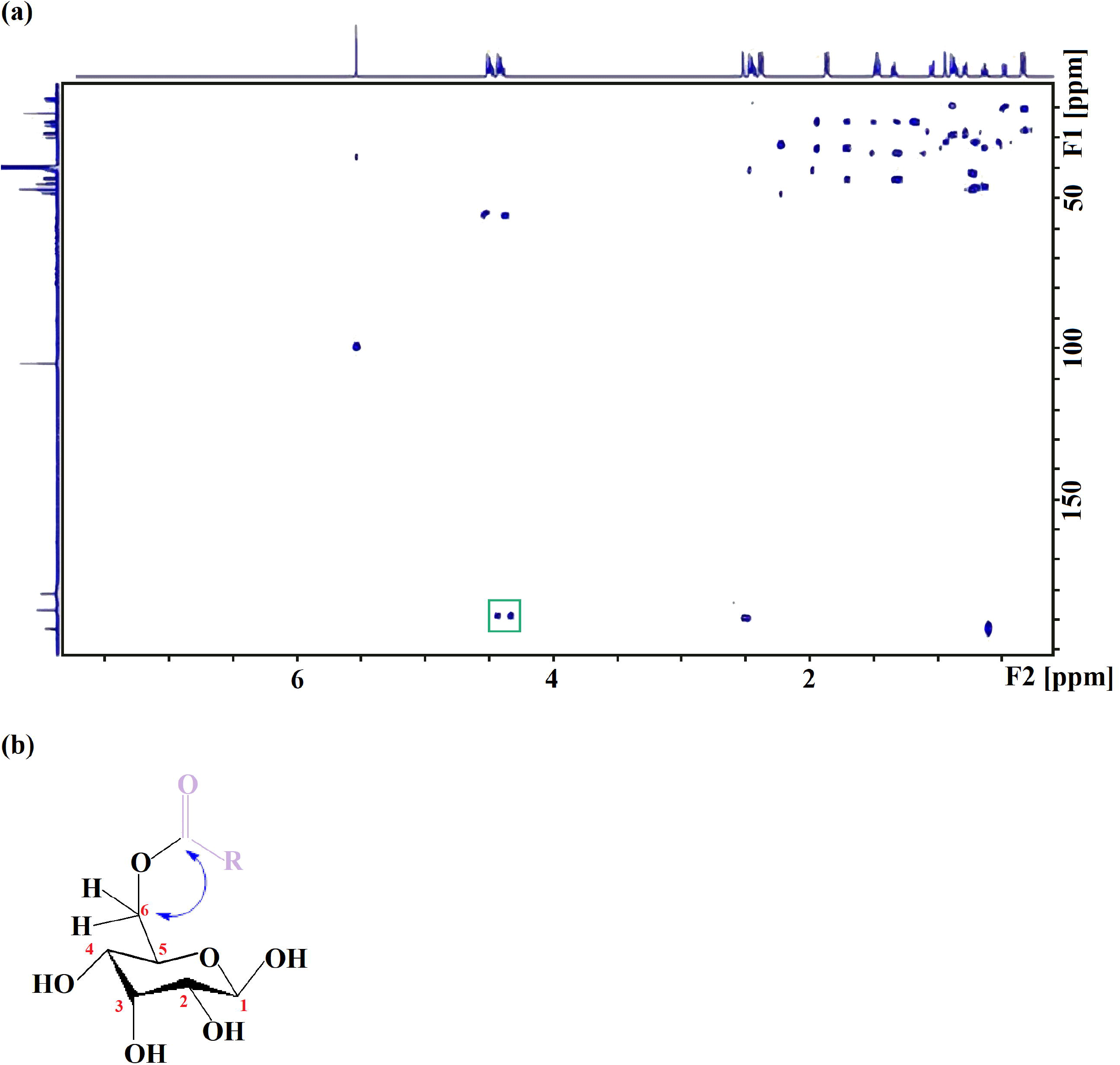
Identification of the ester produced as Glc (C-6)-OCOR by 2D (^1^H-^13^C HMBC) NMR (a) and correlation between the protons of C-6 glucose ester and the carbonyl function in olive oil FAMEs (b).

**Fig. 5.**
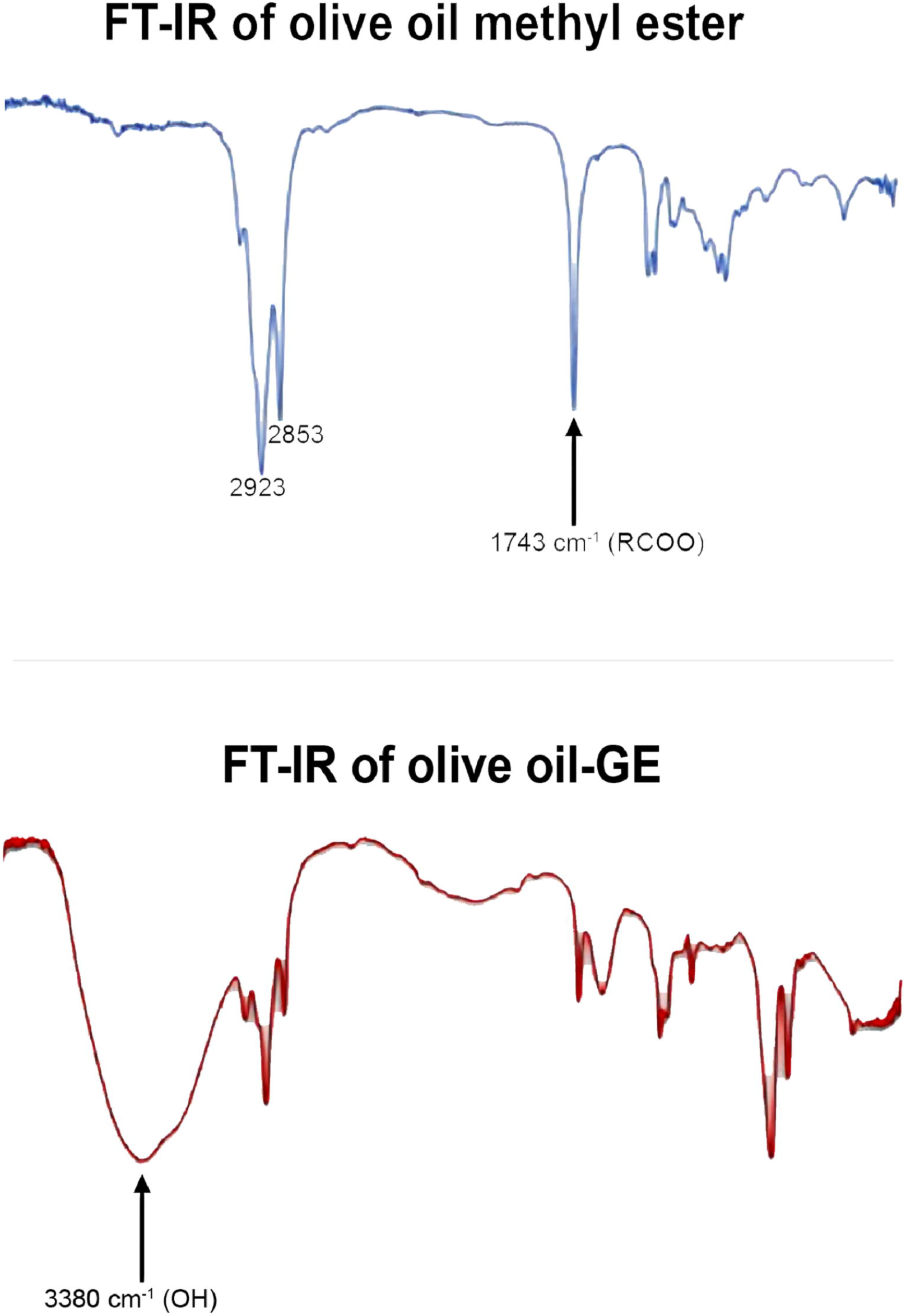
FT-IR of olive oil FAMEs and their glucose esters.

### 3.4. GE synthesis using FAMEs from different origin

After optimization of the reaction conditions GEs were synthesized using FAMEs derived from SCOs produced by *C. echinulata, U. isabellina* and *N. gaditana* and from an EPA concentrate (Table 3). The GE conversion yield when EPA-FAMEs were used as substrate was excellent (i.e. 99%), followed by *N. gaditana-FAMEs, U. isabellina-FAMEs* and *C. echinulata-FAMEs* (i.e. 86, 85 and 80%, respectively – Fig. 6). The structures of the obtained GEs were confirmed on the basis of their FT-IR spectra in which appearance of the broadband due to hydroxyl groups of glucose was observed (Fig. 7 and Fig. S1-S5). According to the above presented data we can conclude that the synthesis of GEs accomplished in this work was successful, while possible large scale applications, using SCOs instead of traditional sources of PUFAs, will not interfere with the food supply chain.

**Fig. 6.**
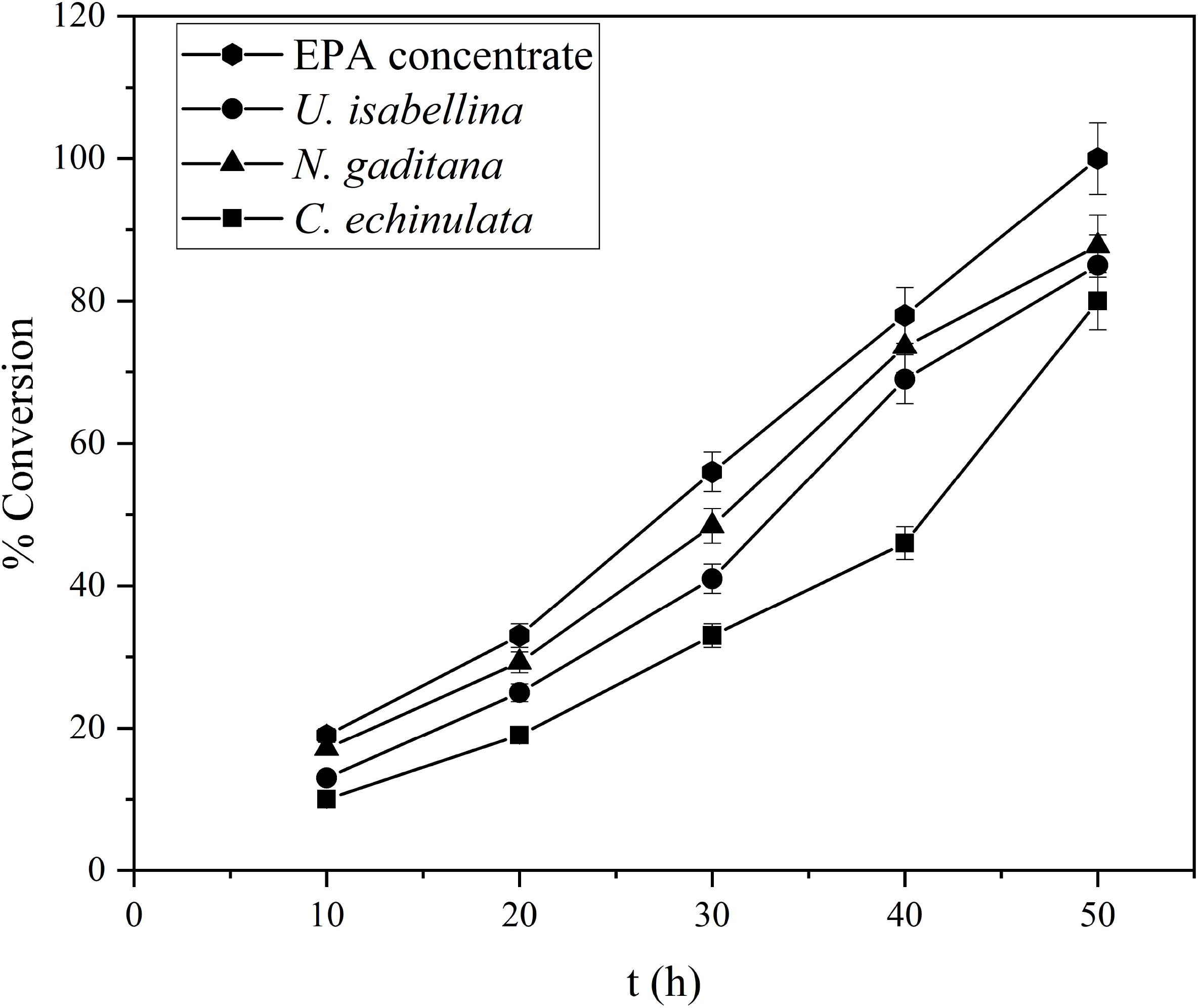
Reaction course over time of conversion of FAMEs derived from different origins to glucose esters.

**Fig. 7.**
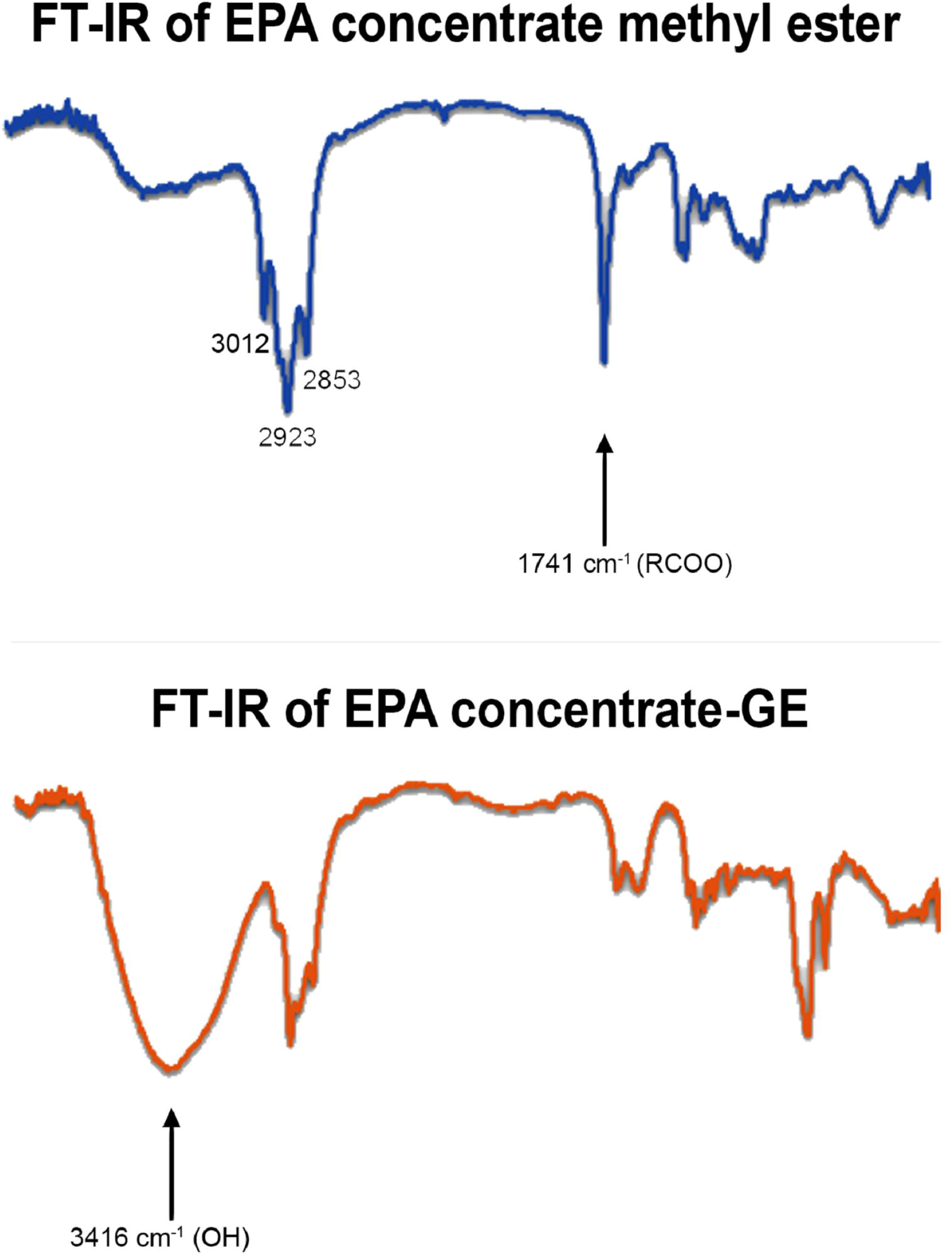
FT-IR analysis of EPA concentrate FAMEs and their glucose esters.

**Table 3.** GE synthesis by lipase CA in a solvent mixture consisted of 80% DMSO and 20% tert-amyl alcohol) utilizing FAMEs of different origin as substrate.

| Entry | Source of FAMES | Time<br>(h) | Conversion<br>(%) |
| --- | --- | --- | --- |
| 1 | *Cunninghamella echinulata* | 24 | 80.3 ± 4.0 |
| 2 | *Umbelopsis isabellina* | 18 | 85.1 ± 4.3 |
| 3 | *Nannochloropsis gaditana* | 18 | 86.3 ± 4.3 |
| 4 | EPA concentrate | 18 | 99.1 ± 5.0 |

### 3.5. Antimicrobial activity of GEs

GEs derived from FAMEs of *C. echinulata, U. isabellina, N. gaditana* SCOs, olive oil and EPA concentrate were tested against various human pathogens for their antimicrobial activity by the agar well diffusion method, which resulted in the formation of a variable diameter zone of inhibition (Table 4).

**Table 4.**
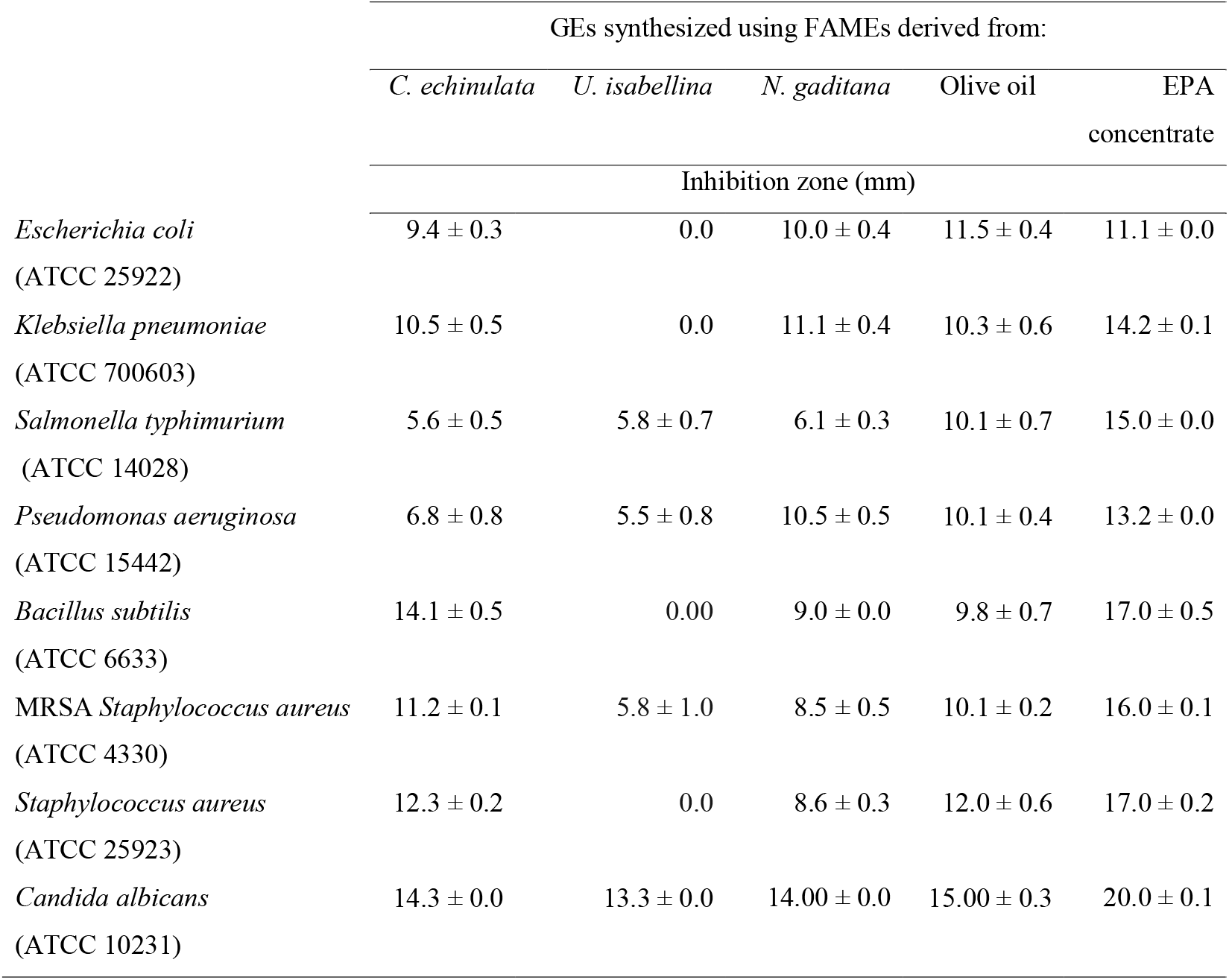
GEs antimicrobial activity using human pathogens as tested organisms. Data represent the mean of three replicates ± SD of the diameter of the inhibition zones.

GEs produced in this work, except for *U. isabellina-GEs*, showed moderate to strong inhibitory activity against all tested organisms. In details, *U. isabellina-GEs* showed weak antimicrobial activity (inhibition zone ≈ 6 mm) against *S. typhimurium, P. aeruginosa* and *S. aureus*. On the contrary, *C. echinulata*-GEs, *N. gaditana-GEs* and olive oil-GEs moderately inhibited all tested organisms, while EPA-GEs showed the strongest antimicrobial activity against all tested organisms, specifically against *C. albicans* (20.00 mm), *B. subtilis* and *S. aureus* (17.00 mm). Finally, all GEs showed a higher antimicrobial activity against *C. albicans* than against bacteria.

It was shown that EPA-GEs were very effective against all tested bacteria and their activity can be attributed to their high EPA content. Furthermore, *C. echinulata-GEs* were more effective against pathogens comparing to *U. isabellina-GEs*, probably due to the presence of GLA in the lipids of *C. echinulata* in enhanced concentrations which is known for its antimicrobial activity [13, 25].

The MIC and MBC values were determined for selected pathogens (Table 5). All tested pathogenic strains are sensitive to all GEs being inhibited at low MIC, ranged between 6.3-50 μg/mL, and destroyed at MBC between 50-100 μg/mL. *C. echinulata-GEs* and EPA-GEs significantly affected the growth of all the tested bacteria than *N. gaditana-GEs* and olive oil-GEs.

**Table 5.** Minimum inhibitory concentration (MIC, μg/mL) and minimum bactericidal concentration (MBC, μg/mL) of the synthesized GEs against pathogens.

| Test organisms | Source of FAMES |  |  |  |  |  |  |  |
| --- | --- | --- | --- | --- | --- | --- | --- | --- |
|  | *C. echinulata* |  | *N. gaditana* |  | Olive oil |  | EPA concentrate oil |  |
|  | MIC | MBC | MIC | MBC | MIC | MBC | MIC | MBC |
| *Klebsilla pneumoniae*<br>(ATCC 700603) | 25.0<br>$\pm 0.0$ | 100.0<br>$\pm 0.0$ | 50.0<br>$\pm 0.0$ | 100.0<br>$\pm 0.0$ | 66.7<br>$\pm 28.8$ | 83.3<br>$\pm 28.8$ | 33.3<br>$\pm 14.4$ | 100.0<br>$\pm 0.0$ |
| *Pseudomonas aeruginosa*<br>(ATCC 15442) | 41.7<br>$\pm 14.4$ | 100.0<br>$\pm 0.0$ | 50.0<br>$\pm 0.0$ | 100.0<br>$\pm 0.0$ | 25.0<br>$\pm 0.0$ | 83.3<br>$\pm 28.8$ | 20.8<br>$\pm 7.2$ | 41.7<br>$\pm 14.4$ |
| *Bacillus subtilis*<br>(ATCC 6633) | 16.7<br>$\pm 7.3$ | 50.0<br>$\pm 0.0$ | 20.8<br>$\pm 7.3$ | 100.0<br>$\pm 0.0$ | 33.3<br>$\pm 14.4$ | 100.0<br>$\pm 0.0$ | 10.4<br>$\pm 3.6$ | 41.7<br>$\pm 14.4$ |
| *Staphylococcus aureus*<br>(ATCC 25923) | 12.5<br>$\pm 0.0$ | 50.0<br>$\pm 0.0$ | 50.0<br>$\pm 0.0$ | 83.3<br>$\pm 28.8$ | 83.3<br>$\pm 28.7$ | 100.0<br>$\pm 0.0$ | 8.3<br>$\pm 3.6$ | 33.3<br>$\pm 14.4$ |

The variability of the inhibitory effect found in this paper is in agreement with previous papers reported that SEs exhibited a variable effect on different bacterial species [52, 53], while depending on the conditions, SEs may inhibit Gram-negative [40, 54] or Gram-positive bacteria [55, 56]. Also, depending on the dose, SEs can be either bactericidal [57] or bacteriostatic [58].

Wagh et al. [59] explained that, the inhibitory effect of SEs is dependent on the esterification level, type (e.g. the length of aliphatic chain) and number of esterified FAs on the sugar molecule and the nature of the carbohydrate. Furthermore, Karlová et al. [60] reported that the antimicrobial effects of fructose esters decreased as the aliphatic chain increased. It seems that the carbon chain length was the most important factor influencing the surface properties, whereas degree of esterification and hydrophilic groups showed little effect [61].

The antimicrobial activity of glucose esters was tested against *E. coli, B. subtilis, B. megaterium* and *B. cereus* [51, 56]. In addition, the unsaturated FAs lactose esters were shown to exhibit antimicrobial activity against Gram-positive, Gram-negative microorganisms and fungi [5]. The antimicrobial activity of SEs is due to autolysis caused by the interaction of the esters with cell membranes of bacteria. The lytic action is thought to be due to the activation of autolytic enzymes rather than the actual solubilization of the bacterial cell membrane [62].

The FAAs synthesized in El-Baz et al. [36] using lipids of similar FA composition as acyl group-donors to those used in this paper, also exhibited a significant antimicrobial activity. However, contrary to the results reported here, the FAAs containing oleic acid in high percentages (i.e. derived from olive oil and *U. isabellina* oil) were more effective against human pathogens than other FAAs.

### 3.6. Insecticidal activity of GEs

*Aedes aegypti* (the yellow fever mosquito) spreads dangerous human arboviruses including dengue, Zika, and chikungunya. Consequently, control of yellow fever mosquitoes is a critical public health priority [63].

The susceptibility of *A. aegypti* larvae to GEs under laboratory conditions was tested using dipping methods. The larvicidal activity of a compound is usually improved by increasing its concentration and exposure time, as Rodrigues et al. [64] reported for plant-derived bioactive products, such as essential oils, ethanol extracts and FAMEs. In the current study, *C. echinulata*-GEs showed a strong insecticidal activity against *A. aegypti* larvae with LC50 0.541 mg/L, which could be probably attributed to the presence of GLA in significant concentrations, followed by EPA-GEs, olive oil-GEs, and *N. gaditana-GEs*, demonstrating LC50 10.24, 12.88 and 16.92 mg/L, respectively. Contrary, *U. isabellina-GEs* were less active, presenting a LC50 equal to 39.62 mg/L (Table 6, Fig. 8). On the other hand, the RR values indicated that mosquito of *A. aegypti* was much more susceptible to *C. echinulata-GEs* than to EPA-GEs, olive oil-GEs, *N. gaditana-GEs* and *U. isabellina-GEs* by about 20 to 70 folds. Overall, most of the GEs produced in this study, especially those of *C. echinulata-GEs*, had superior insecticidal activity. Likewise, FAAs synthesized using GLA rich lipids produced by *C. echinulata*, displayed a superior insecticidal activity against the same organism [36], suggesting that this FA is probably a key molecule, responsible for the bioactivity of preparations.

**Fig. 8.**
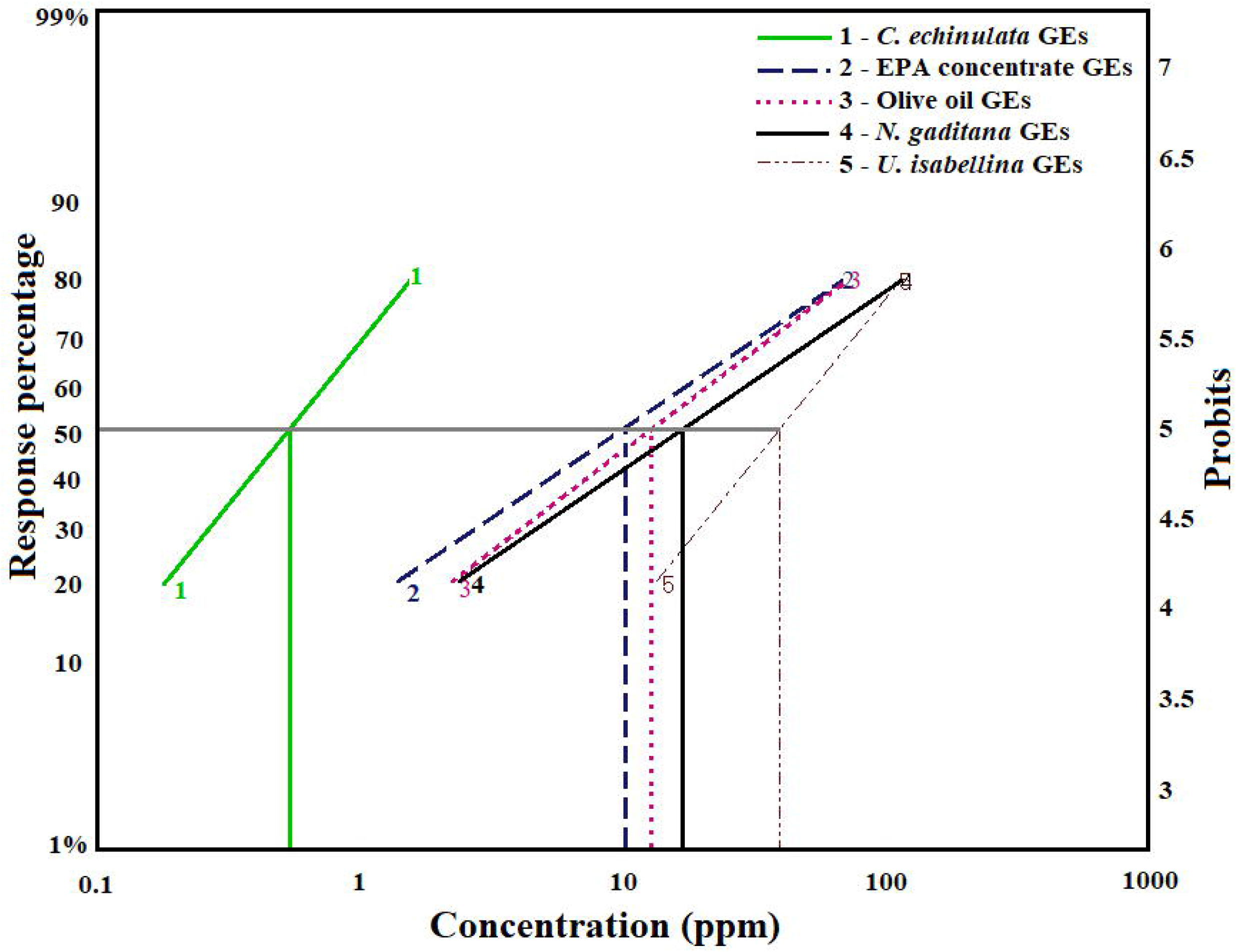
The larval mortality effect of GEs derived from *C. echinulata*, EPA concentrate, olive oil, *U. isabellina* and *N. gaditana* at different concentrations against *Aedes aegypti* after continuous exposure for 48 h.

**Table 6.**
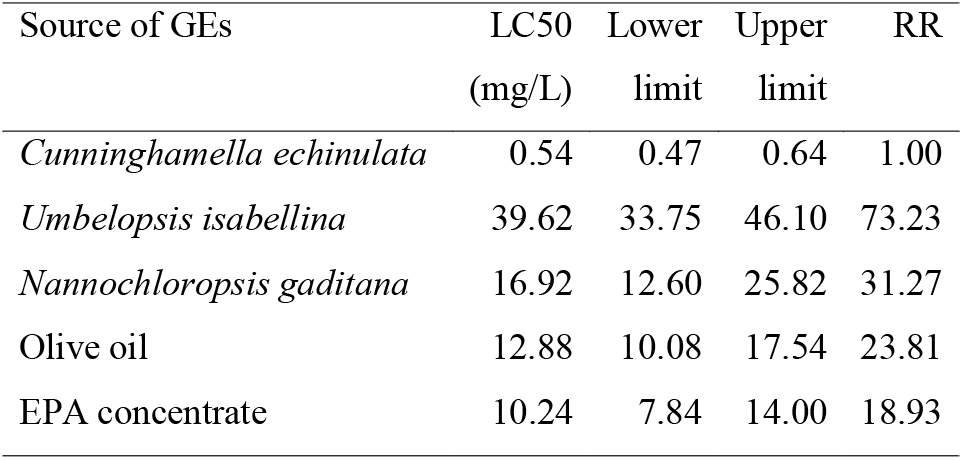
Susceptibility of *Aedes aegypti* larvae to GEs under laboratory conditions by using dipping methods. Data represent the mean values of six replicates.

| Source of GEs | LC50<br>(mg/L) | Lower<br>limit | Upper<br>limit | RR |
| --- | --- | --- | --- | --- |
| *Cunninghamella echinulata* | 0.54 | 0.47 | 0.64 | 1.00 |
| *Umbelopsis isabellina* | 39.62 | 33.75 | 46.10 | 73.23 |
| *Nannochloropsis gaditana* | 16.92 | 12.60 | 25.82 | 31.27 |
| Olive oil | 12.88 | 10.08 | 17.54 | 23.81 |
| EPA concentrate | 10.24 | 7.84 | 14.00 | 18.93 |

Chortyk [65] reported that SEs are useful as effective, environmentally safe pesticides for the control of soft-bodied arthropod pests. In addition, Puterka et al. [6] confirmed that the majority of the SEs exhibited higher insecticidal activity than insecticide soap. The nature of both the sugar and FA moieties determine the SE physicochemical properties, such as the solubility in water and stability of emulsions, and their insecticidal activity. However, changing the sugar or FA components from lower to higher carbon chains or whether the sugar is a monosaccharide or disaccharide does not follow a consistent relationship to insecticide activity. C18 FAs, such as oleic, elaidic, linoleic, and linoleic acids inhibited proliferation of malarial parasites in mice infected with *Plasmodium vinckei petteri* or with *Plasmodium yoelii nigeriensis* [66].

### 3.7. Quantitative analysis of ovarian cancer cell apoptosis induced by GEs

The ability of both FAMEs and GEs produced in this study to induce SKOV-3 cell apoptosis was assessed by flow cytometry after Annexin FITC staining of cells (Fig. 9). The results show that all GEs induced apoptosis of the SKOV-3 ovarian cancer cell line compared to untreated cells, with the apoptotic rate increasing significantly after 48 h. A higher percentage of apoptosis was observed in the cells treated with EPA-GEs (i.e. 43.1%), followed by *C. echinulata-GEs, U. isabellina-GEs* and olive oil-GEs (i.e. 39.2, 34.0 and 33.5%, respectively). Similarly, FAAs containing EPA in their structures in high percentages displayed a strong anticancer activity against SKOV-3 ovarian cancer cell line [36].

**Fig. 9.**
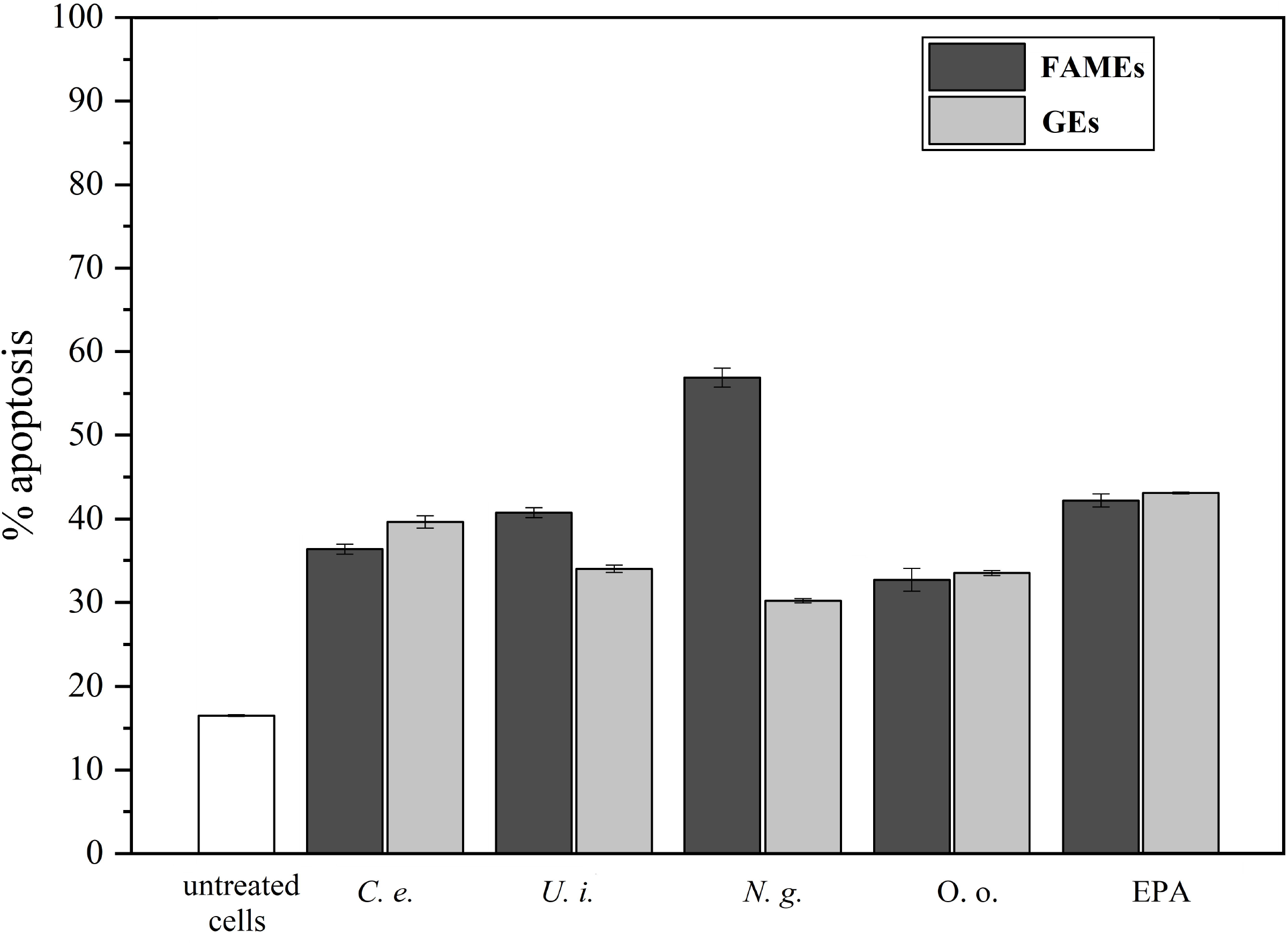
Effect of FAMEs and GEs on SKOV-3 cancer cell line. Flow cytometry analysis of apoptosis in SKOV-3 cells either untreated or treated with 10 μg/mL of every compound for 48 h. After the treatment period, the cells were stained with Annexin FITC and subsequently analyzed by flow cytometry.

Morin et al. [67] and Siena et al. [68] reported that, a variety of modified FAs are promising molecules in the treatment of cancers. Also, there were several studies dealing with anticancer, antimicrobial and anti-inflammatory activities of SE derivatives [69, 70]. Our results correlated with those reported by An and Feng [71], whose evaluated the antitumor activity of series of glucosyl esters derivatives against three cancer cells, human breast adenocarcinoma (MCF-7), human colon carcinoma (K562) and human hepatoma (HepG2). They found that the glucosyl esters exhibited significant anticancer activity in dose- and time-dependent fashion. The structure - activity relationship analysis revealed that lipophilic properties might be an essential parameter affecting their activity. Research to inhibit cancer cell proliferation has shown that SE activity is linked to the nature of both sugar and fatty acyl chain [71].

### 4. Conclusions

Two immobilized lipases, especially lipase CA, efficiently catalyzed the synthesis of SEs using glucose and FAMEs, derived from lipids of different origin including SCOs, as substrates. The reaction of GE synthesis can be completed under environmentally friendly conditions using a solvent mixture consisted of 80% DMSO and 20% tert-amyl alcohol in 24 h. The enzyme used in the synthesis can be recycled at least three times without losing its catalytic activity. The synthesized GEs displayed significant biological activities against important human pathogenic microorganisms, the larvae of *A. aegypti* and the SKOV-3 ovarian cancer cell line, which are related to their FA profile. Hence, we can conclude that SCOs, characterized by a wide diversity in FA composition, can be considered as sources of acyl group-donors suitable for the production of GEs of different bioactivity.

## Author agreement

Hatim A. El-Baz, Ahmed M. Elazzazy, Tamer S. Saleh, Marianna Dourou, Jazem A. Mahyoub, Mohammed N. Baeshen, Hekmat R. Madian and George Aggelis have all agreed to submission.

## Acknowledgment

This work was funded by the *University of Jeddah*, Saudi Arabia, under grant No. (UJ-06-18-ICP). The authors, therefore, acknowledge with thanks the University technical and financial support.

## Abbreviations

ANOVA: Analysis of variance
EPA: eicosapentaenoic acid
FAs: fatty acids
FAAs: fatty acid amides
FAMEs: fatty acid methyl esters
FT-IR: Fourier-transform infrared spectroscopy
FFAs: free fatty acids
GEs: glucose fatty acid esters
GLA: γ-linolenic acid
HMBC: heteronuclear multiple bond correlation
Lipase CA: lipase from *Candida antarctica*
Lipase CR: lipase from *Candida rugosa*
MBC: minimum bactericidal concentration
MIC: minimum inhibitory concentration
NMR: Nuclear magnetic resonance
PUFAs: polyunsaturated fatty acids
SCOs: single cell oils
SEs: sugar esters
TLC: thin-layer chromatography

## Abbreviations

C. e.: C. echinulata
U. i.: U. isabellina
N. g.: N. gaditana
O. o.: Olive oil
EPA: EPA concentrate

