## Supplementary tables and figures for "Singe cell oil (SCO) based bioactive compounds: II. Enzymatic synthesis of glucose fatty acid esters using SCOs as acyl group-donors and their biological activities": Supplementary material.docx

**Table S1** Biomass yield (x, g or mg/L) and lipid content (L/x, %) of the microorganisms used in this study as source of lipids. The cultures were performed in triplicate.

| **Microorganism** | **x** | **L/x (%)** |
| --- | --- | --- |
| *Cunninghamella echinulata* | 12.9 ± 0.9 g/L | 30.0 ± 1.5 |
| *Umbelopsis isabellina* | 13.2 ± 1.2 g/L | 74.0 ± 0.8 |
| *Nannochloropsis gaditana* | 313.9 ± 0.4 mg/L | 22.7 ± 0.1 |

**Table S2** Fatty acid composition of the methyl ester mixtures used as acyl-donors in the amide and GEs synthesis. Analyses were performed in three independent samples.

| Sample | Fatty acid composition (%, w/w) | | | | | | | | | | |
| --- | --- | --- | --- | --- | --- | --- | --- | --- | --- | --- | --- |
|  | C16:0 | C16:1  n-7 | C18:0 | C18:1  n-9 | C18:2  n-6 | C18:3 n-6 | C18:3 n-3 | C18:4  n-3 | C20:1  n-9 | C20:5  n-3 | Others |
| *Cunninghamella echinulata* | 15.9  ± 0.7 | 2.0  ± 0.5 | 7.9  ± 0.6 | 44.0  ± 1.4 | 13.0  ± 1.2 | 12.8  ± 1.0 | nd | nd | nd | nd | 4.4  ± 1.2 |
| *Umbelopsis isabellina* | 22.1  ± 1.1 | 3.6  ± 0.4 | 2.8  ± 0.4 | 54.4  ± 4.1 | 11.7  ± 0.9 | 2.6  ± 0.3 | nd | nd | nd | nd | 2.8  ± 0.7 |
| *Nannochloropsis gaditana* | 18.0  ± 0.7 | 20.4  ± 0.5 | 0.7  ± 0.0 | 13.7  ± 0.4 | 5.7  ± 0.1 | 0.8  ± 0.2 | 0.8  ± 0.1 | nd | 7.2  ± 0.4 | 25.0  ± 0.3 | 7.7  ± 2.7 |
| Olive oil | 12.2  ± 1.2 | 2.4  ± 0.2 | 2.7  ± 0.3 | 74.1  ± 1.1 | 7.0  ± 0.2 | nd | 0.4  ± 0.0 | nd | nd | nd | 1.2  ± 0.3 |
| EPA concentrate | 0.5  ± 0.0 | 0.5  ± 0.0 | 3.3  ± 0.2 | 10.0  ± 1.8 | 1.0  ± 0.1 | 0.8  ± 0.0 | 2.3  ± 0.2 | 2.4  ± 0.1 | 4.4  ± 0.8 | 72.3  ± 1.4 | 4.0  ± 0.5 |

The original spectra of FT-IR analysis of *Cunninghamella echinulata*, *Umbelopsis isabellina*,

*Nannochloropsis gaditana*, Olive oil and EPA concentrate FAMEs and their glucose esters are presented in the Figs. S1-S5.

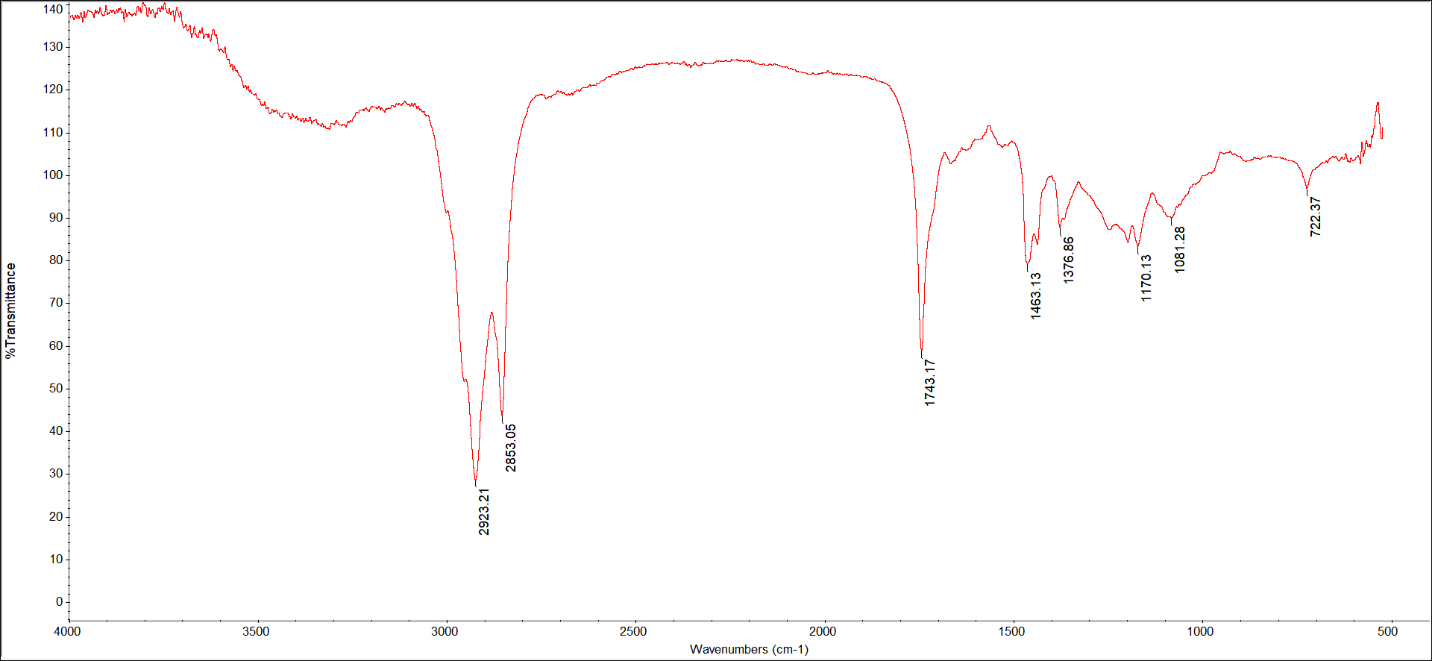

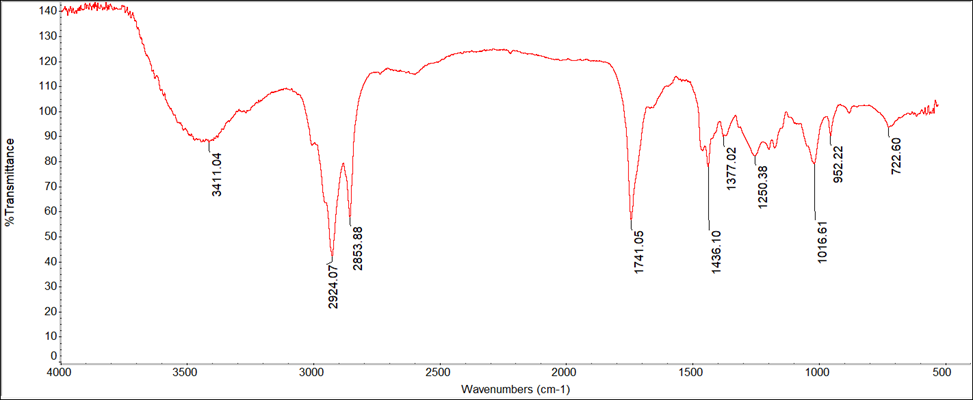

**Fig. S1** FT-IR analysis of *Cunninghamella echinulata* FAMEs and their glucose esters.

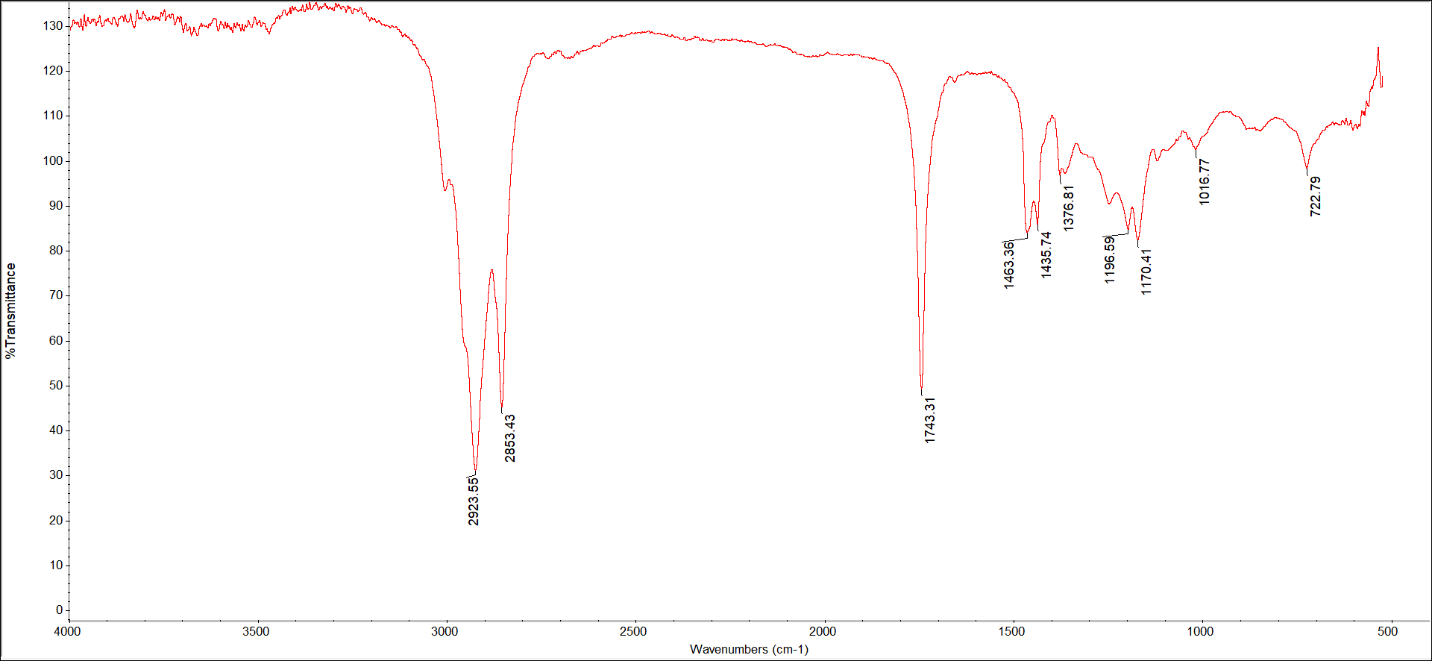

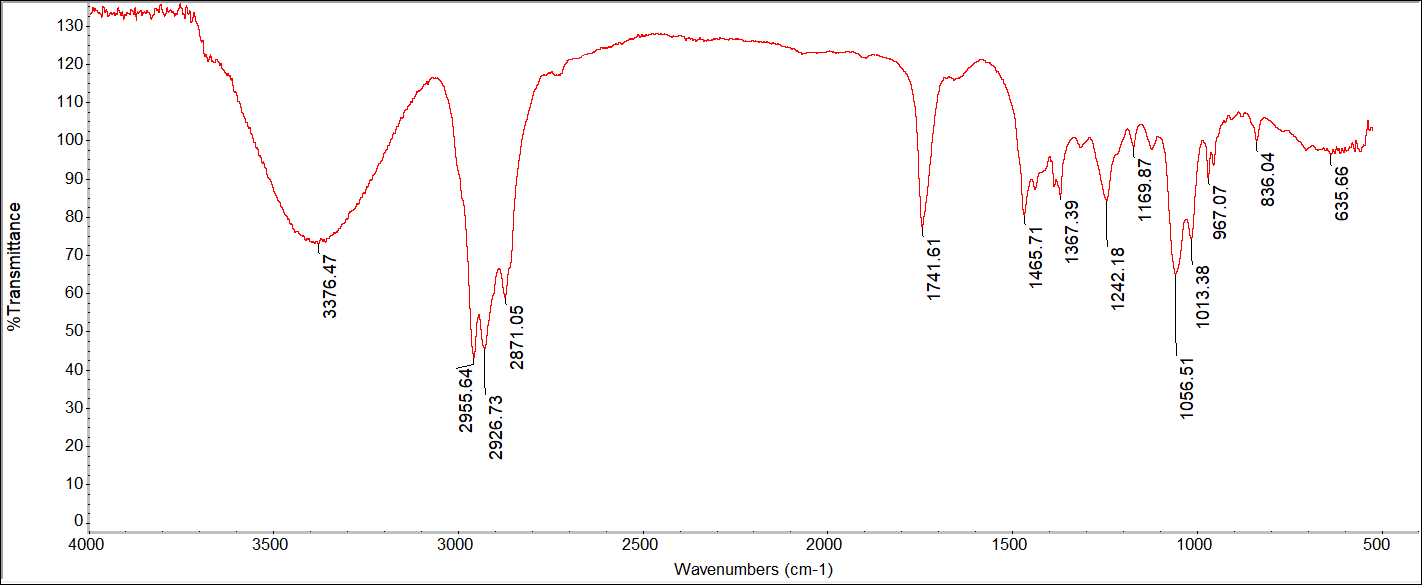

**Fig. S2** FT-IR analysis of *Umbelopsis isabellina* FAMEs and their glucose esters.

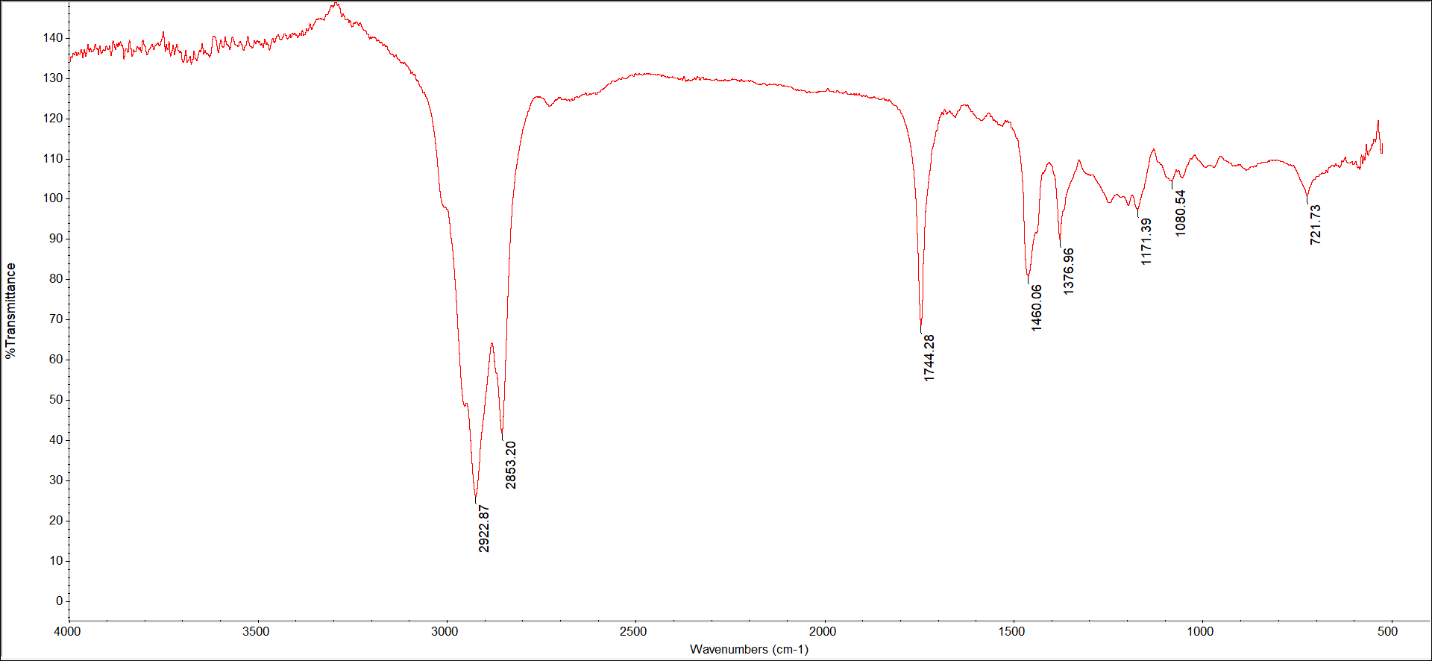

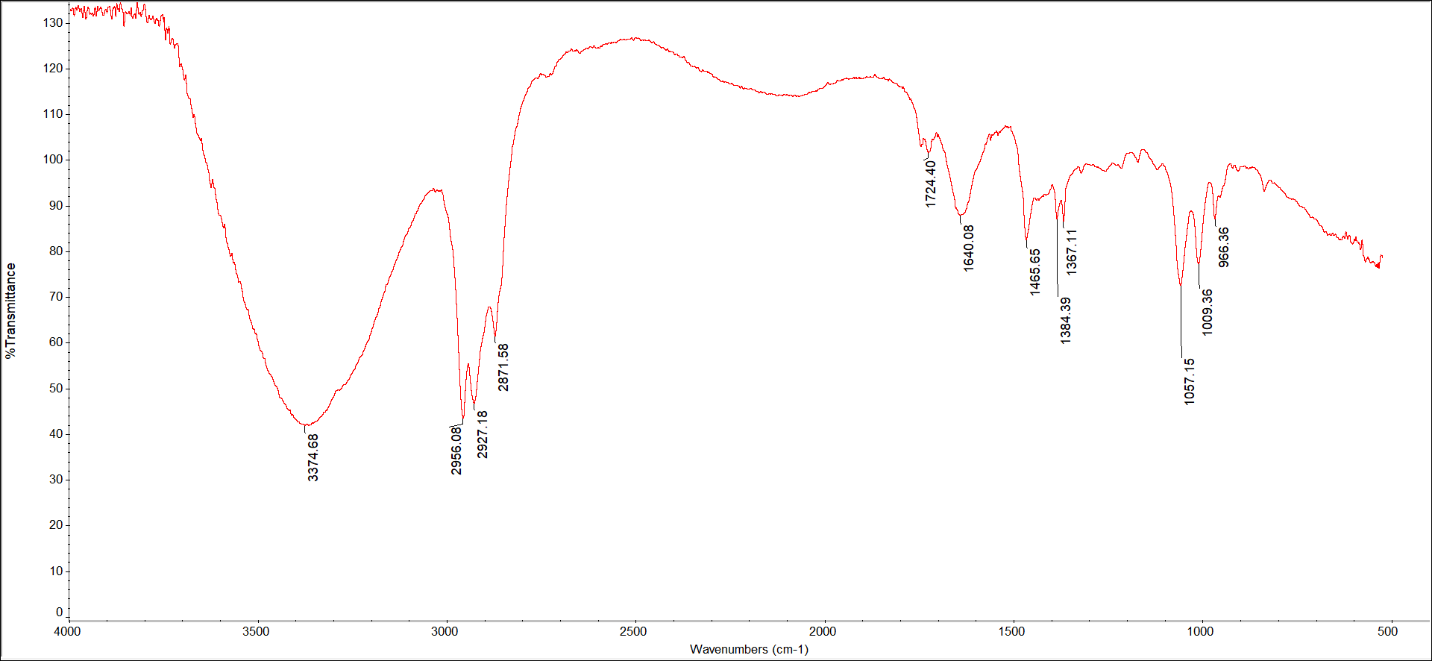

**Fig. S3** FT-IR analysis of *Nannochloropsis gaditana* FAMEs and their glucose esters.

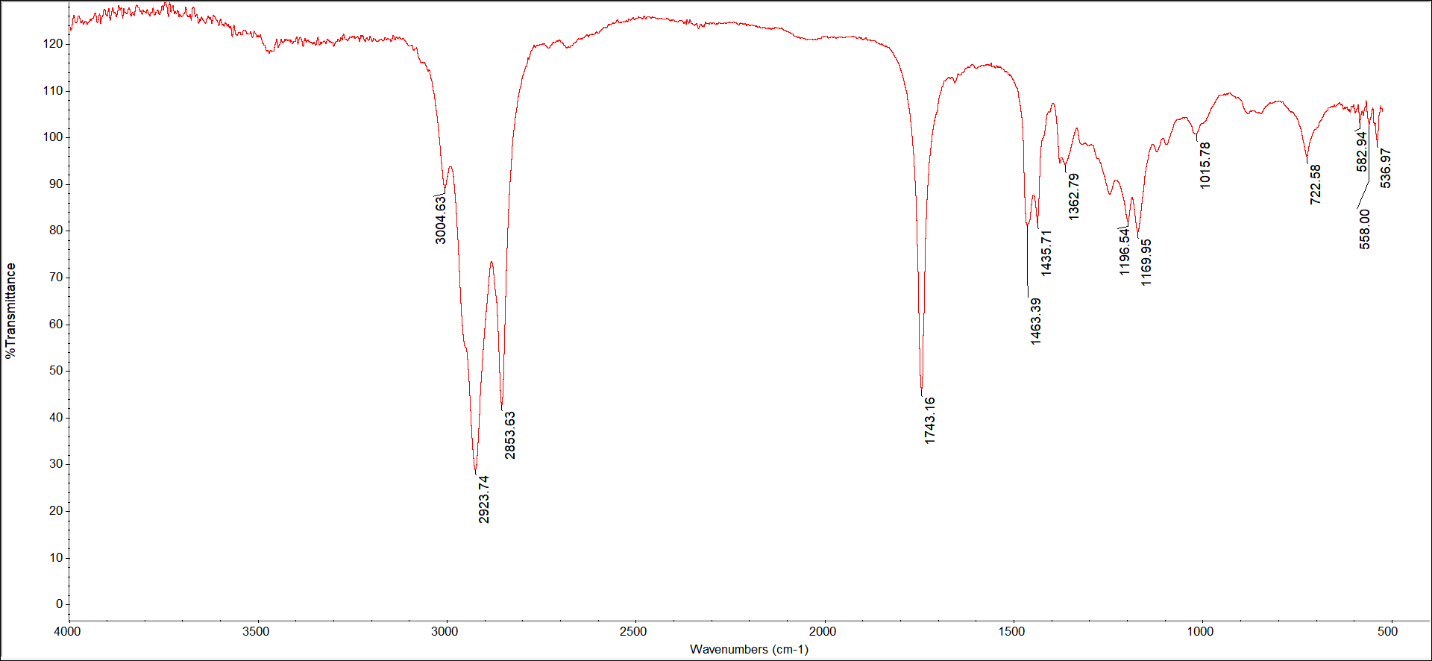

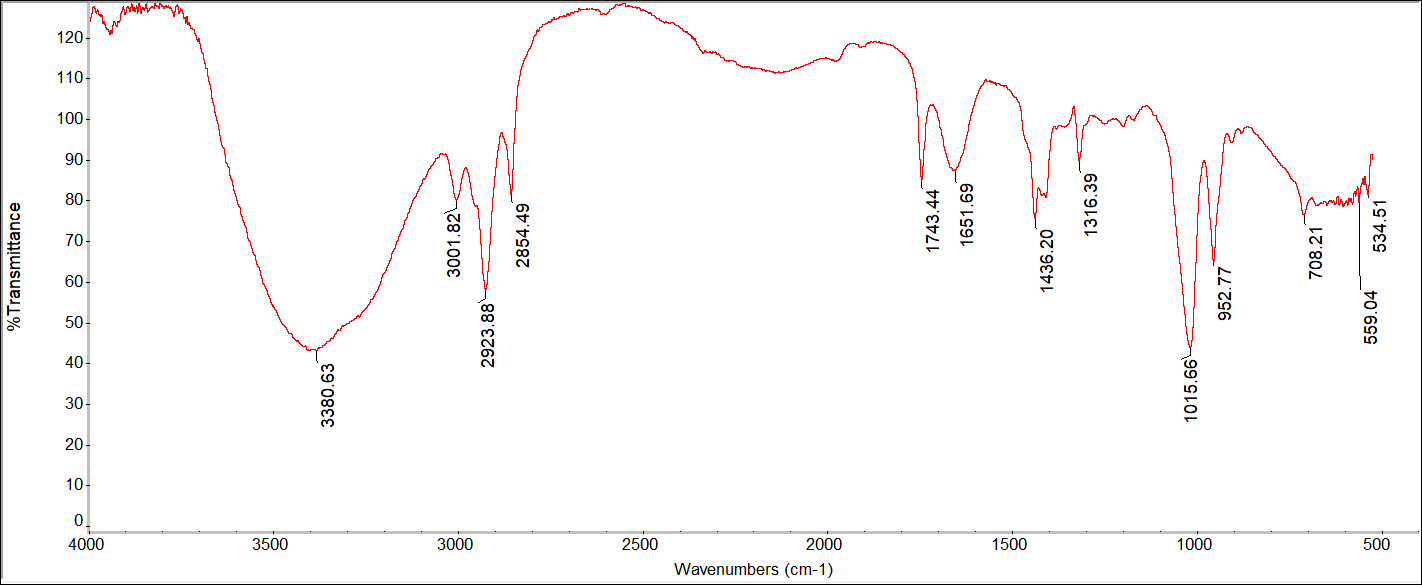

**Fig. S4** FT-IR analysis of olive oil FAMEs and their glucose esters.

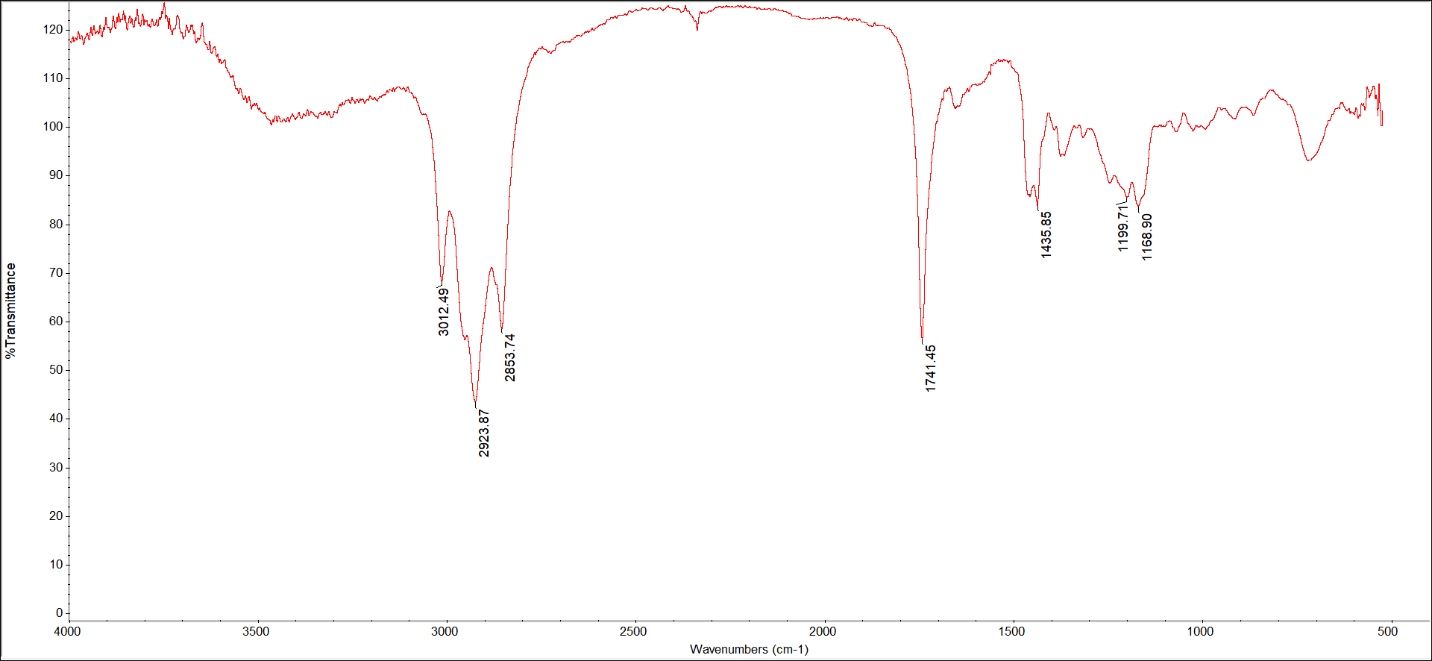

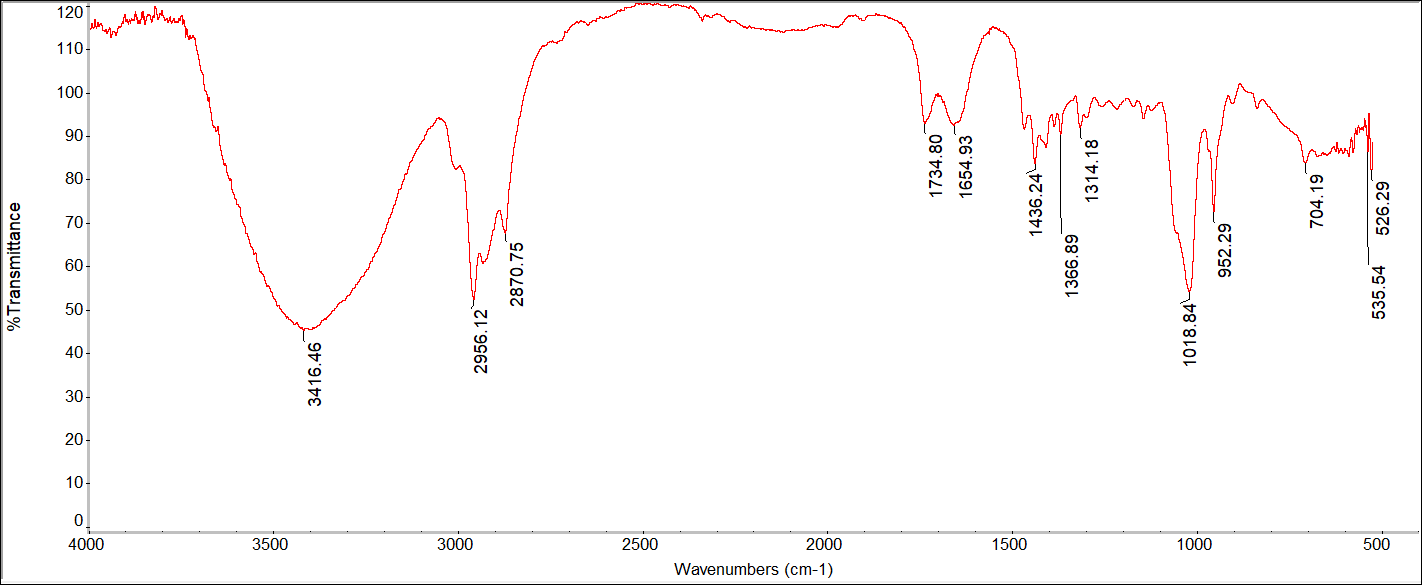

**Fig. S5** FT-IR analysis of EPA concentrate FAMEs and their glucose esters.
